# Differential targeting of human pyroptotic Caspase-5 and Caspase-4 by *Shigella* OspC2 and OspC3

**DOI:** 10.64898/2025.12.04.692159

**Authors:** Kyungsub Kim, Marcos Antonio Valdespino Diaz, Rowan Karr, Michele E. Muscolo, Lisa Goers, Truelian Yu, Tera C. Levin, Jonathan N. Pruneda, Cammie F. Lesser

## Abstract

Pyroptosis is an inflammatory cell death pathway that is a key defense mechanism of intestinal epithelial cells. To successfully establish an infection, intracytosolic Gram-negative pathogens must block this host response. Indeed, *Shigella* OspC3 suppresses epithelial pyroptosis by targeting and inactivating Caspase-4 (CASP4). Here, we demonstrate that OspC2, which shares 96% identity with OspC3, targets Caspase-5 (CASP5), a close paralog of CASP4. Through a combination of yeast two-hybrid, transfection, and bacterial infection assays, we show that the distinct pyroptotic caspase specificities of OspC2 and OspC3 are determined by a short α-helical region, designated the <u>P</u>yroptotic <u>C</u>aspase <u>S</u>pecificity (PCS) domain. This domain is located upstream from the <u>A</u>nkyrin <u>R</u>ich <u>R</u>epeat (ARR) region previously established to promote OspC3 binding to CASP4. Swapping PCS domains between OspC2 and OspC3 is sufficient to redirect their caspase targeting. Evidence for CASP5-driven pyroptosis in response to infection has not yet been established. However, CASP5 displays signatures of positive selection at residues predicted to interact with the PCS domain of OspC2. Notably, the introduction of orangutan-specific residues into human CASP5 disrupt its interaction with and modification by OspC2, demonstrating that CASP5 natural variation can cause key functional differences in this host-microbe molecular interaction. These findings highlight the evolutionary interplay between bacterial effectors and host proteins, support a likely role for CASP5 in responding to Gram-negative bacteria, and identify promising therapeutic targets for enhancing epithelial defense against bacterial pathogens.

**Importance:** *Shigella* species are human-specific Gram-negative pathogens that establish a replicative niche in intestinal epithelial cells by blocking pyroptosis, a key inflammatory cell death pathway. We reveal that two closely related *Shigella* type III secreted effectors, OspC2 and OspC3, specifically inactivate the human caspase paralogs CASP5 and CASP4, respectively. This specificity is determined by their newly identified Pyroptotic Caspase Specificity (PCS) domains. In addition, we find that positively selected residues in CASP5 alter the OspC2/CASP5 interaction, underscoring the impacts of ongoing evolutionary arms races between bacterial effectors and host immune proteins. By elucidating the molecular basis of caspase targeting and adaptation, this work provides new insight into the diversification of host defense mechanisms and identifies potential therapeutic targets for enhancing epithelial resistance to bacterial infection.

## Introduction

Pyroptosis is an inflammatory form of cell death that plays a critical role in the innate immune response to bacterial infection^1^. Pyroptosis is triggered upon sensing bacterial components, including lipopolysaccharide (LPS), a major constituent of the outer membrane of Gram-negative bacteria. Pyroptosis is mediated via the activation of pyroptotic caspases, including CASP1, CASP4, CASP5, and CASP11^2^. Once activated these proteases promote the processing of gasdermin D (GSDMD) and proinflammatory cytokines, including IL-18 and IL-1β. The N-terminal fragment of GSDMD proceeds to forms pores in the host cell plasma membrane that enable the release of the pro-inflammatory cytokines and the eventual death of the infected cells via a lytic pathway^3,4^.

The intestinal epithelium serves as the first line of defense against invading microbes. While canonical CASP1 and non-canonical CASP11 inflammasomes play major roles in mounting responses to enteric pathogens in mouse intestinal epithelial cells^5,6^, CASP1 is minimally expressed in human intestinal epithelial cells^5,7,8^. Within these cells, non-canonical inflammasomes are the main pyroptotic signaling pathway activated, primarily via activation of CASP4^9,10^. Similar to CASP4, its close paralog CASP5, is activated upon binding LPS^11,12^. However, while CASP5 has been found to cooperate with CASP4 in responding to invading pathogens within macrophages^13,14^, a role for it in inducing pyroptosis in epithelial cells has not yet been established.

*Shigella* species, the causative agents of bacterial dysentery (shigellosis), are model intracytosolic bacterial pathogens. Humans are the natural hosts of *Shigella,* although non-human primates are also susceptible to infection. In macrophages, *Shigella* uptake rapidly induces cell death primarily via the activation of canonical CASP1 inflammasomes, resulting in the release of the *Shigella* from the dying cells^15,16^. In contrast, upon entering the cytosol of intestinal epithelial cells, *Shigella* trigger the activation of non-canonical CASP4 inflammasomes^17^. However, inflammasome activity is blocked via the activity of multiple injected type III effectors^9,10,17–22^ and *Shigella* establishes a replicative niche within the colonic epithelium.

The virulence of *Shigella*, like that of many other Gram-negative bacteria, is dependent on a type III secretion system (T3SS). *Shigella* uses this nanomachine to translocate∼30 distinct proteins^23,24^, referred to as effectors, directly into the cytosol of host cells^25^. One of these effectors, OspC3, is the founding member of a family of bacterial effectors that ADP-riboxanate host proteins^19,26,27^. *Shigella* encodes three OspC effectors^23^, OspC1, OspC2 and OspC3. OspC2 and OspC3 share 96% sequence identity with each other and 63% with OspC1. These proteins have both shared and unique host cell targets. In terms of caspases, OspC3 has been established to target CASP4^17,19^ and OspC1 to modify CASP8. All three OspC effectors also modify protein components of the eukaryotic translation initiation 3 (eIF3) complex^30^.

Here, using complementary yeast two-hybrid, transfection, and bacterial infection assays, we have established that OspC2 and OspC3 selectively bind and inactivate CASP5 and CASP4, respectively. We found that this specificity is defined by a short α-helical region, the <u>P</u>yroptotic <u>C</u>aspase <u>S</u>pecificity (PCS) domain, which is located upstream from the <u>A</u>nkyrin <u>R</u>ich <u>R</u>epeat (ARR) region previously established to promote OspC3 binding to CASP4. Indeed, swapping the PCS domains of OspC2 and OspC3 is sufficient to switch the caspase targeting specificity of each effector. Furthermore, we found that the PCS domain of OspC2 likely interacts with some of the rapidly evolving residues in CASP5 and that this CASP5 natural variation altered OspC2 targeting. These studies provide an example of how effectors have evolved to precisely target specific host proteins, reflecting the ongoing molecular arms race between bacteria and the host innate immune system.

## Results

### mT3Sf, a bottom-up approach to study *Shigella* type III secreted effectors

The roles of type III secreted effectors in pathogenesis are typically studied by screening for phenotypes associated with their absence. To complement this approach, we previously developed a DH10β *E. coli-*based bottom-up platform (mT3.1 *E coli*), referred to here as mT3Ec to screen for phenotypes associated with their presence^20,21^. The operons encoding the components needed to form the *Shigella* type III secretion apparatus (T3SA) and almost all of its effectors are encoded on a large virulence plasmid^23^. mT3Ec is a non-pathogenic strain that contains a smaller plasmid engineered to carry the three operons (ipa, mxi, and spa) that encode the components that form the T3SA plus 4 effectors (IpaA, IpgB1, IcsB and IpgD) plus a second plasmid that encodes their shared transcriptional regulator, VirB, expressed under the control of the PTRC, an IPTG (Isopropyl-β-D thiogalactopyranoside)-inducible promoter^20^.

We previously established that mT3Ec invades and escapes into the cytosol of infected epithelial cells triggering their cell death via pyroptosis, which we found in our first bottom-up screen was suppressed by the addition of several effectors^20^. However, given potential limitations of this strain, as DH10β *E. coli* lacks *Shigella’s* chromosomally-encoded pathogenicity islands, and O-antigen chains on its LPS^31^, we developed mT3Sf, a *Shigella-*based variant (Fig. 1A). The chassis in this case is BS103, a virulence-plasmid minus *Shigella* strain^32^. A notable difference between mT3Ec and mT3Sf is that in the latter case, the *ipa, mxi,* and *spa* operons were introduced onto the chromosome. Furthermore, to generate a version of mT3Sf that does not express or secrete any of *Shigella’s* chromosomally-encoded effectors, we deleted the gene encoding MxiE, their shared transcriptional regulator^24,33,34^, from the *mxi* operon. Variants of mT3Sf that carry the VirB expression plasmid secrete components of the T3SA tip complex (IpaB, IpaC, and IpaD) at levels only slightly lower than *S. flexneri*(Fig. 1B). All the subsequent mT3Sf experiments described herein, were conducted with the Δ*mxiE* variant of mT3Sf.

**Fig. 1.**
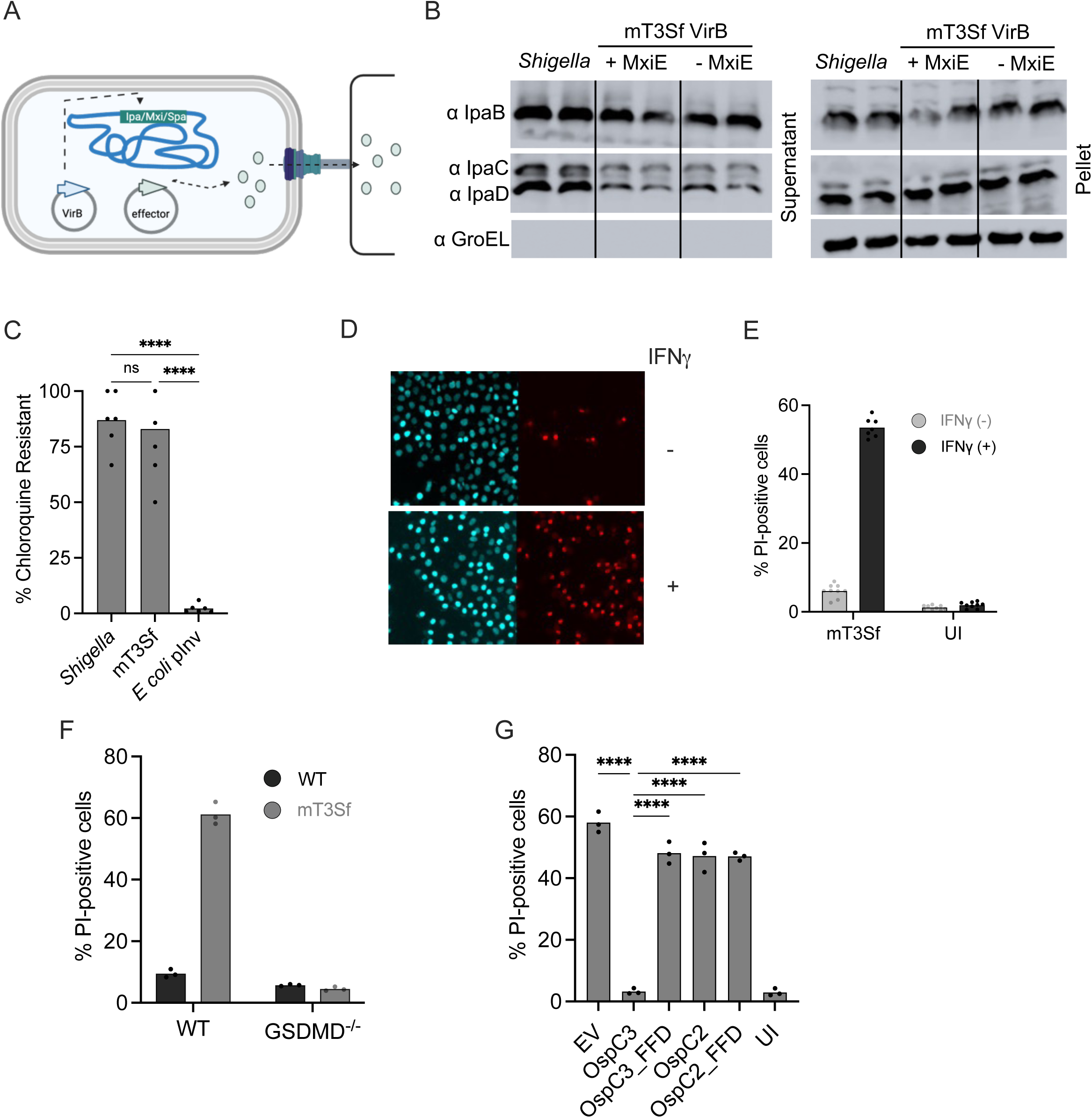
Infection with mT3Sf triggers pyroptosis of epithelial cells. (A) Schematic of mT3Sf. (B) Secretion assays comparing the levels of IpaB, IpaC and IpaD secreted by WT *Shigella* and mT3Sf VirB +/- MxiE. Cell equivalent levels of secreted (supernatant) and bacterial (pellet) fractions were analyzed. GroEL serves as a lysis/loading control. Blot is representative of two experimental repeats. (C) CHQ^R^ assay: HeLa cells were infected with designated strain at an MOIs (100 for *Shigella,* 10 for mT3Sf and *E. coli* expressing *Yersinia* Invasin) for 1 h and then treated with gentamicin (gent) ± CHQ for an additional 1 h before cell monolayers were lysed and bacteria enumerated. The percentage of chloroquine resistant bacteria = (CHQ/Gent)^R^/Gent^R^ bacteria. Infections were conducted in triplicate. (D-H) HeLa cells unprimed (D-E) or primed with IFNγ (D-G) were infected with the designated mT3Sf strains at an MOI of 5. One-hour post-infection, gentamicin, PI and Hoechst were added to the media. After an additional 2 hours, the cells were imaged (D) and the percentage of PI+ cells quantified (E-G). Each infection condition included at least three technical repeats and that shown is representative of at least 3 independent assays. Statistical significance when indicated was assessed by one-way ANOVA with Tukey’s post hoc test. *P < 0.05, **P < 0.01, ****P < 0.0001, ns = nonsignificant.

As previously observed for mT3Ec, we found that mT3Sf can invade and escape into the cytosol of epithelial cells. This is not surprising, as three of the four effectors encoded with the *ipa* and *mxi* operons (IpgB1, IpaA, and IpgD) promote the invasion of *Shigella* into epithelial cells^35^. We found that essentially all intracellular gentamicin-resistant mT3Sf and *Shigella* were resistant to chloroquine, an antimicrobial, which only accumulates to levels high enough to kill intraphagosomal bacteria. In contrast, almost all gentamicin-resistant intracellular *E. coli* that encode *Yersinia* Invasin (Inv), an adhesin that promotes their uptake into phagosomes^36^, were killed when infected cells were treated with chloroquine (Fig. 1C).

### OspC3 but not OspC2 suppresses mT3Sf-triggered cell death

Given that mT3Sf invade and escape into the cytosol of epithelial cells, we reasoned that their invasion into host cells would trigger cell death via pyroptosis. To investigate the fate of the infected cells, we used an automated microscopy-based assay, whereby infected cells were treated with Hoescht and propidium iodide (PI), membrane-permeable and membrane-impermeable dyes, respectively. PI is a marker of pyroptosis as it can enter cells through GSDMD pores. As we previously observed with mT3Ec, infection with mT3Sf, unlike WT *Shigella,* induced epithelial cell death, as indicated by an increased detection of PI+ cells, which was enhanced when the cells were primed with IFNγ^20,21^ (Fig. 1D-E). We observed minimal cell death of mT3Sf infected GSDMD^-/-^ epithelial cells (Fig. 1F). Together, these observations strongly suggest that infection with mT3Sf triggers epithelial cell death via pyroptosis.

*Shigella* OspC2 shares 96% identity with OspC3, yet only OspC3 binds and post-translationally modifies CASP4^19,26^. We previously observed that infection with mT3Ec::OspC3 triggered minimal cell death of infected epithelial cells. Similarly, we found that IFNγ-primed epithelial cells infected with mT3Sf:EV, but not mT3Sf::OspC3, were highly likely to undergo cell death. In contrast, we found that cells infected with mT3Sf::OspC2, as well as with variants of mT3Sf that secrete catalytically-dead (FFD: F188A, F211A, D231A) variants of OspC2 and OspC3^26^ (Fig. S1A), were as likely to undergo cell death as those infected with mT3Sf::EV (Fig. 1G). Furthermore, infection with mT3Sf triggered minimal cell death of IFNγ-primed CASP4^-/-^ cells (Fig. S2A-B). Together, these observations suggested that infection with mT3Sf primarily triggers cell death via activation of CASP4 inflammasomes.

### OspC2 and OspC3 bind differently to human pyroptotic caspases

Given the high degree of similarity of OspC2 with OspC3 and between CASP4 and CASP5, we hypothesized that OspC2 targets CASP5. CASP4 and CASP5 share a high degree of homology with mouse CASP11 and arose via a primate-specific duplication^37^. All three of these pyroptotic caspases are composed of three domains (Fig. 2A): an N-terminal CARD (caspase recruitment domain) followed by large (L) and small (S) protease subunits. Upon the sensing of LPS via CARD, the self-cleaved, liberated L and S subunits assemble into an active cysteine protease that cleave GSDMD and pro-inflammatory cytokines.

**Fig. 2.**
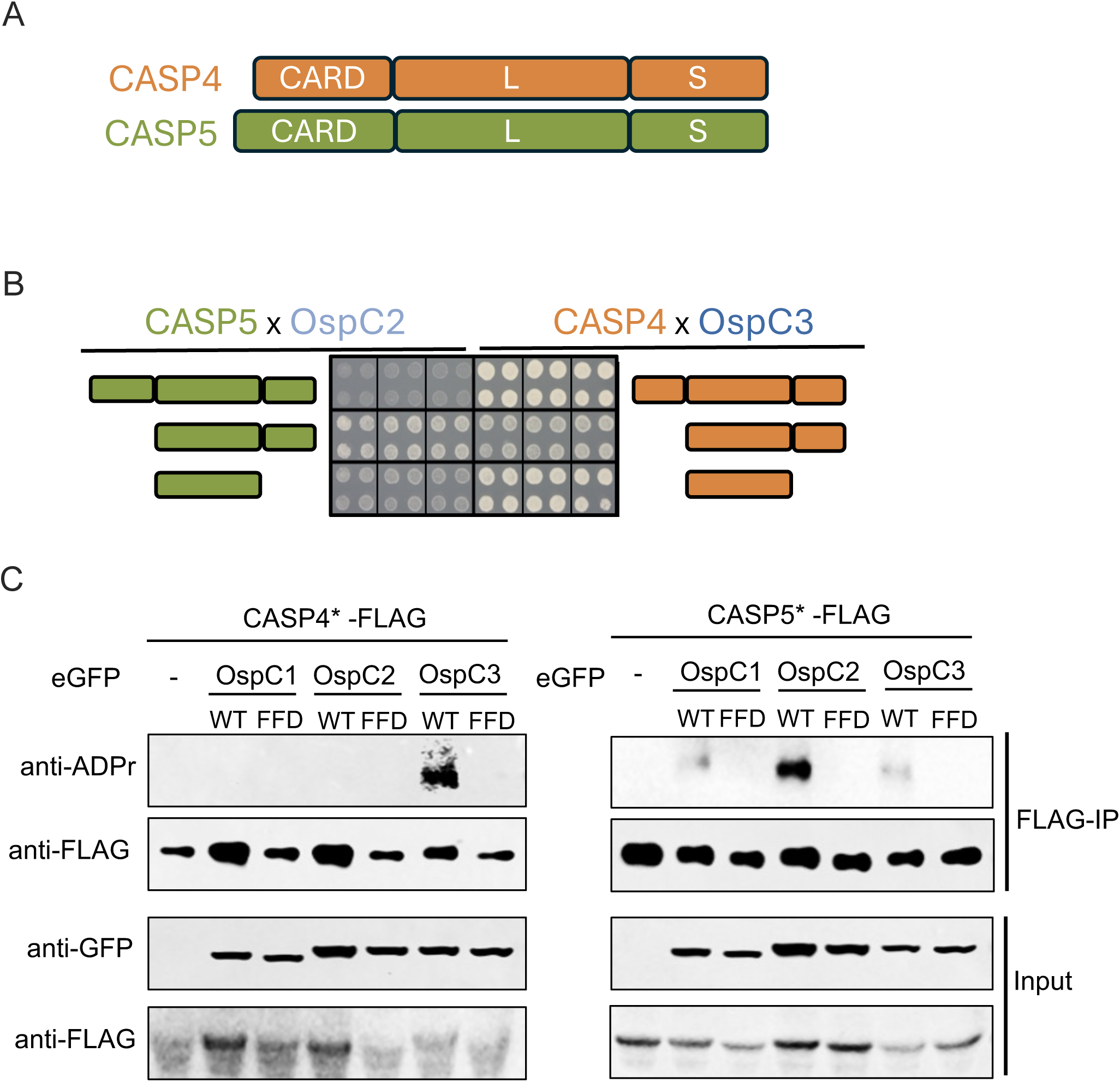
OspC2 and OspC3 distinctly bind and modify CASP5 and CASP4, respectively. (A) Schematic of the domain compositions of CASP4 and CASP5. (B) Images of the Y2H interactions detected between OspC2/CASP5 and OspC3/CASP4 obtained after 7 days of growth on selective media. (C) HEK293 cells were co-transfected with plasmids that express the designated GFP-OspC effector plus one that expresses a 3xFLAG tagged catalytically dead full-length variant of CASP4 or CASP5. After 24h, the cells were lysed and the FLAG-tagged caspases were immunoprecipitated. Immunoblots of the input and immunoprecipitated fractions were probed with the designated antibodies. The blots shown are representative of at least three experimental repeats.

Several lines of experimentation, including co-immunoprecipitation assays^17^ and a genome-wide yeast-two hybrid (Y2H) screen^38^ have established that OspC3 can bind to the L and L-S fragments of CASP4. In addition, a co-crystal of OspC3 in complex with the L-S CASP4 fragment has been solved^26^. Given the previous Y2H findings, we investigated whether we could detect a similar Y2H interaction between OspC2 and CASP5. Thus, we compared the ability of OspC2 and OspC3 to bind to the full-length CASP4 and CASP5 as well as their L and L-S fragments. As expected, OspC3 interacted with all three variants of CASP4 (Fig. 2B). In addition, excitingly, OspC2 interacted with both the L and L-S fragments of CASP5 (Fig. 2B). This interactions were specific, as neither OspC2 nor OspC3 interacted with any additional full length human caspases1-10 (Fig. S3A) or their L domains (Fig. S3B). Furthermore, only OspC3 bound to the L-S fragment of CASP4 (Fig. S3C). These studies provided the first evidence to suggest that OspC2 targets CASP5.

### OspC2 targets the post-translational ADP-riboxanation of CASP5

We next investigated whether we could monitor the ADP-riboxination activity of the *Shigella* OspC effectors. As shown by others^19,30^ we found that numerous host cell proteins were detected on an immunoblot of transiently transfected HEK cells that express WT but not catalytically-dead FFD/AAA variants of GFP-OspC1, GFP-OspC2 and GFP-OspC3^30^ probed with an antibody that recognizes ADP-riboxination modifications (Fig. S4A).

To assess the ability of each OspC effector to specifically modify CASP4 or CASP5, we co-transfected the HEK cells with each WT and catalytically dead GFP-effector plus a FLAG-tagged caspase variant, 24 hours later, we immunoprecipitated the caspases and investigated whether they were modified. As expected, WT but not catalytically dead OspC3 specifically modified CASP4 (Fig. 2C). Similarly, we found that WT, but not catalytically-dead, OspC2 uniquely modified CASP5 (Fig. 2C). OspC1 did not modify either caspase (Fig. 2C). Interestingly, as previously observed for OspC3, we found that OspC2-mediated modification of CASP5 was blocked in the presence of ZVAD, a small molecule inhibitor, which binds to its active site. This observation suggests that OspC2, like OspC3^26^, modifies substrate-free caspases (Fig. S4B). These data provided the first evidence that OspC2 and OspC3 differentially modify CASP5 and CASP4.

### Determination of the OspC2 and OspC3 regions responsible for caspase binding specificity

OspC2 and OspC3 are each composed of 454 amino acids organized into three domains (Fig. 3A). Their first ∼50 N-terminal amino acids contain an N-terminal type III secretion signal sequence followed by a chaperone binding domain^39^, their middle ∼300 amino acids encode their catalytic ADP-riboxanation domain and calmodulin-binding regions^26,38^, and their C-terminal ∼100 amino acids, (which contain several ankyrin rich repeats (ARRs), form their ankyrin repeat domain (ARD). In the case of OspC3, the ARD was established to promote its binding to CASP4^17,26,40^. The ARDs of OspC2 and OspC3 differ at 14 of 108 positions (indicated by vertical lines in Fig. 3B), only two of which (demarcated in red in Fig. 3B) were previously established to mediate the binding of OspC3 to CASP4^26^. Based on this observation, we hypothesized that additional amino acids in the ARDs of OspC2 and OspC3 define their pyroptotic caspase binding specificities.

**Fig. 3.**
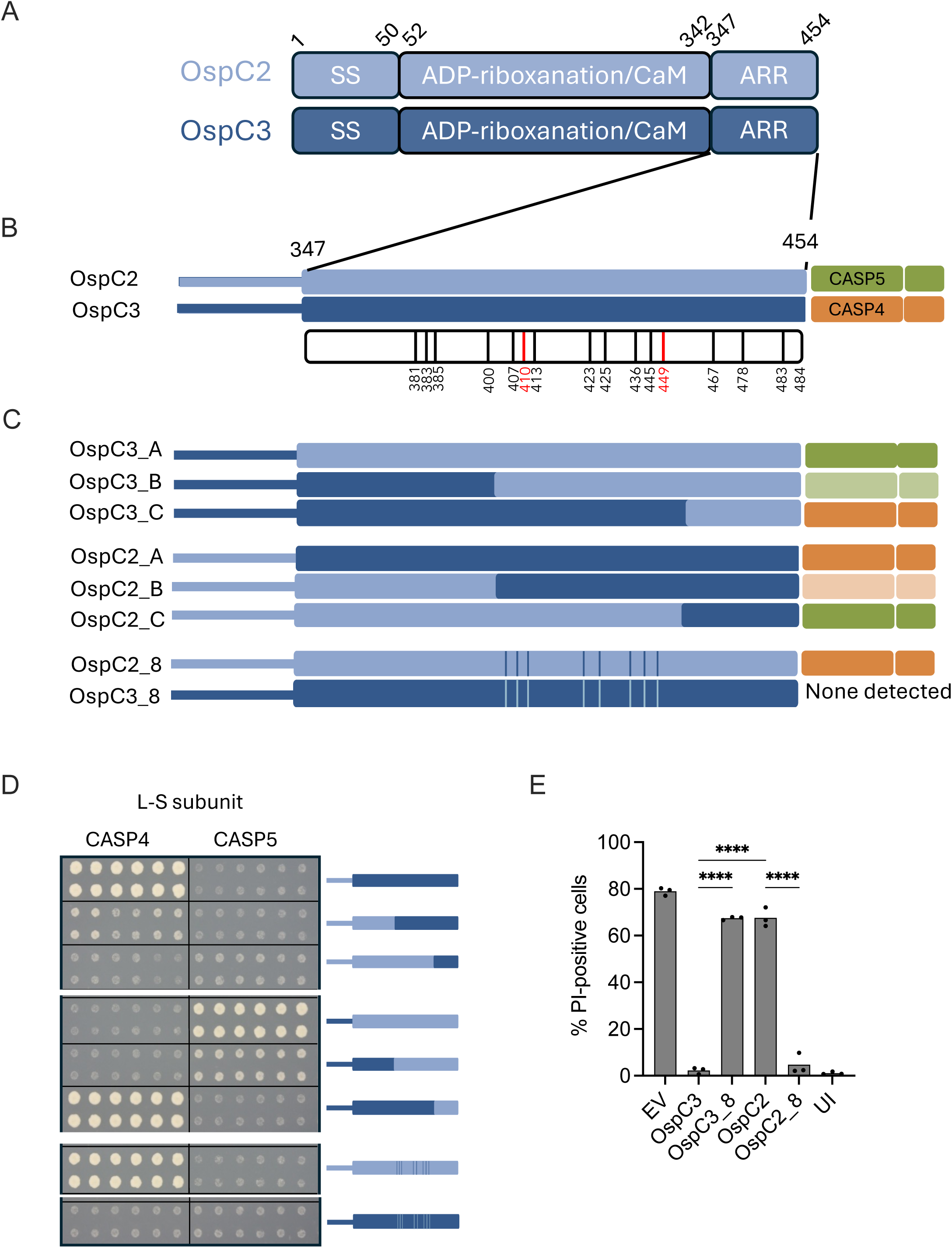
OspC2 and OspC3 differentially recognize CASP4/CASP5 via a defined region within their ARDs. (A) Schematic of the domain compositions of OspC2 and OspC3. (B-D) Summary of the Y2H interactions detected between OspC2/C3 chimeras with the L-S fragments of CASP4 and CASP5. The numbered bars in B represent positions of differing amino acid between OspC2 and OspC3. The red bars indicate residues were previously established to promote OspC3-CASP4 binding. (E) IFNγ-primed HeLa cells were infected with the designated mT3Sf strains at an MOI of 5. One-hour later, gentamicin, PI and Hoechst were added to the media. After Two hours, the cells were imaged and the percentage of PI+ cells quantified. Each infection condition included at least three technical repeats and that shown is representative of at least 3 independent assays. Statistical significance when indicated was assessed by one-way ANOVA with Tukey’s post hoc test. *P < 0.05, **P < 0.01, ****P < 0.0001, ns = nonsignificant.

To narrow down the amino acids that define their caspase binding specificities, we generated a set of reciprocal OspC2/OspC3 chimeras, focusing on their ARDs (Fig. 3C). Using the Y2H assay, we assessed the binding of each to the L-S fragments of CASP4 and CASP5 (Fig. 3C-D). When the entire ARDs or the C-terminal 80 amino acids were swapped, the binding specificity of each chimera matched that of the C-terminal region from which it was derived. Specifically, OspC3_A and OspC3_B interacted with CASP5, while OspC2_A and OspC2_B interacted with CASP4. In contrast, when the C-terminal ∼20 amino acids were swapped, the binding specificity mirrored that of the N-terminal region from which it originated. OspC3_C interacted with CASP4, while OspC2_C interacted with CASP5. While we were unable to assess binding to full-length CASP5, we similarly found that OspC2_A, OspC2_B, and OspC3_C, but not OspC3_A, OspC3_B, and OspC2_C, interacted with full-length CASP4 (Fig. S5).

These observations suggested that the region conferring caspase-binding specificity is located between amino acids 407 and 449, the outer boundaries of the B and C swaps. Indeed, introducing amino acids 407-449 of OspC3 into OspC2 resulted in OspC2_8, a variant that binds to both full-length CASP4 (Fig. S5) and its L-S fragment (Fig. 3C-D). The reciprocal OspC3-based variant, OspC3_8, did not bind to either CASP5 or CASP4. This could be because the binding affinity of OspC3_8 and CASP5 is below the threshold of detection in the Y2H assay or due to impaired stability of OspC3_8 in yeast.

To test the physiological relevance of the Y2H findings, we compared the fate of IFNγ-primed epithelial cells infected with mT3Sf::OspC3, mT3Sf::OspC2, mT3Sf::OspC3_8, mT3Sf::OspC2_8, and mT3Sf::EV. Cells infected with mT3Sf::OspC3 and mT3Sf::OspC2_8 exhibited essentially no evidence of cell death, while those infected with mT3Sf::OspC2 and mT3Sf::OspC3_8 exhibited similar levels of PI-uptake as those infected with mT3Sf::EV (Fig. 3E). Each of the OspC variants was secreted at equivalent levels (Fig. S1B). Together, these observations strongly suggest that OspC2_8, like OspC3, targets and inhibits the activity of CASP4.

### OspC effector family members that interact with pyroptotic caspases possess a distinctive α-helix

Building upon our chimera mapping experiments, we sought to precisely identify the amino acid residues of OspC2 and OspC3 responsible for their differential targeting of CASP5 and CASP4. The chimera-based studies indicated that at least some of the eight differing amino acids, located at positions 407, 410, 413, 423, 425, 436, 445, and 449, contribute to their distinct binding specificities (Fig. 4A). Interestingly, three of these residues (407, 410, and 413) are positioned on a short α-helix, located just upstream of the ARR region within the ARD of OspC3, which was previously identified as crucial for its interaction with CASP4^19^. Notably, in the solved OspC3-CASP4 co-crystal structure, this α-helix is positioned in close proximity to CASP4^26^ (Fig. 4B).

**Fig. 4.**
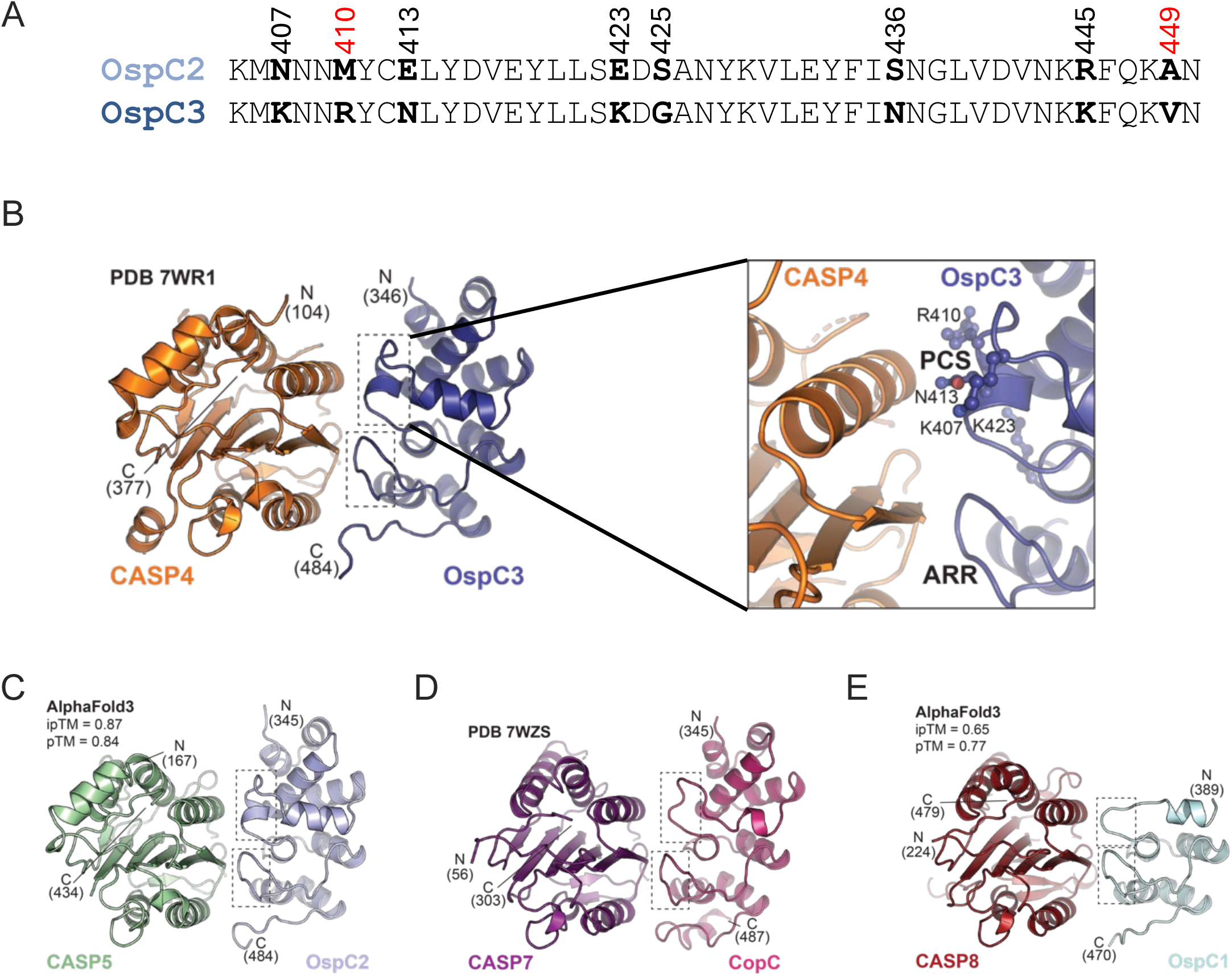
The effector region interacting with pyroptotic caspase possesses a distinct α-helical domain. (A) Alignment of amino acid residues located between positions 405 and 450 of OspC2 and OspC3. Number residues are those that differ in the two effectors. Numbers shown in red, represent residues of OspC3 previously established to promote binding to CASP4. (B-E) 3D crystal structures of the L-S fragment of designed caspases in complex with the ARRs of labeled complexes. In each panel, dotted boxes are drawn around the regions corresponding to the PCS and ARR domains of OspC3. The inlet shows blow of region containing OspC3 PCS domain (C-E). The images in (B, D) are representative of published crystal structures and (C, E) are AlphaFold3 models.

Strikingly, the structural prediction generated by AlphaFold 3 for OspC2 in complex with CASP5 revealed a comparable α-helix in OspC2 positioned opposite CASP5 (Fig. 4C). Furthermore, this α-helix is absent from the solved structure of *Chromobacterium violaceum* CopC in complex with CASP7^37^ (Fig. 4D) and the AlphaFold 3-predicted structure of *Shigella* OspC1 in complex with CASP8 (Fig. 4E). As CASP7 and CASP8 are apoptotic caspases, this α-helix was unique to the OspC family members that target pyroptotic caspases. Interestingly, a sequence alignment of the ARDs of >30 OspC homologs revealed that OspC1 and OspC2, and the homolog from *Escherichia fergusonii,* a potential opportunistic pathogen^41^, were found to likely encode this α-helix (Fig. S6A-B). Together, these observations led us to hypothesize that this short α-helix is critical in defining the binding specificities of OspC2 and OspC3 for CASP4 and CASP5.

### Identification of the OspC <u>P</u>yroptotic <u>C</u>aspase <u>S</u>pecificity (PCS) Domain

We next sought to identify the minimal set of OspC3-specific amino acids that, when introduced into OspC2, would be sufficient to generate a variant that when present in mT3Sf suppresses cell death of infected cells. This experimental approach was predicated on the assumption that OspC2 variants capable of suppressing cell death do so by modifying and inactivating CASP4. Our initial focus was on residues 407, 410, and 413, all located within the OspC2/3-specific α-helix. Introducing these three residues from OspC3 into OspC2 resulted in a variant (OspC2_3A), which partially suppressed mT3Sf-induced cell death (Fig. 5A). Substitution of only two residues (410 and 413; OspC2_2) or a single residue (410; OspC2_1) led to progressively diminished suppression, highlighting the significance of these positions within the helix (Fig. 4B).

**Fig. 5.**
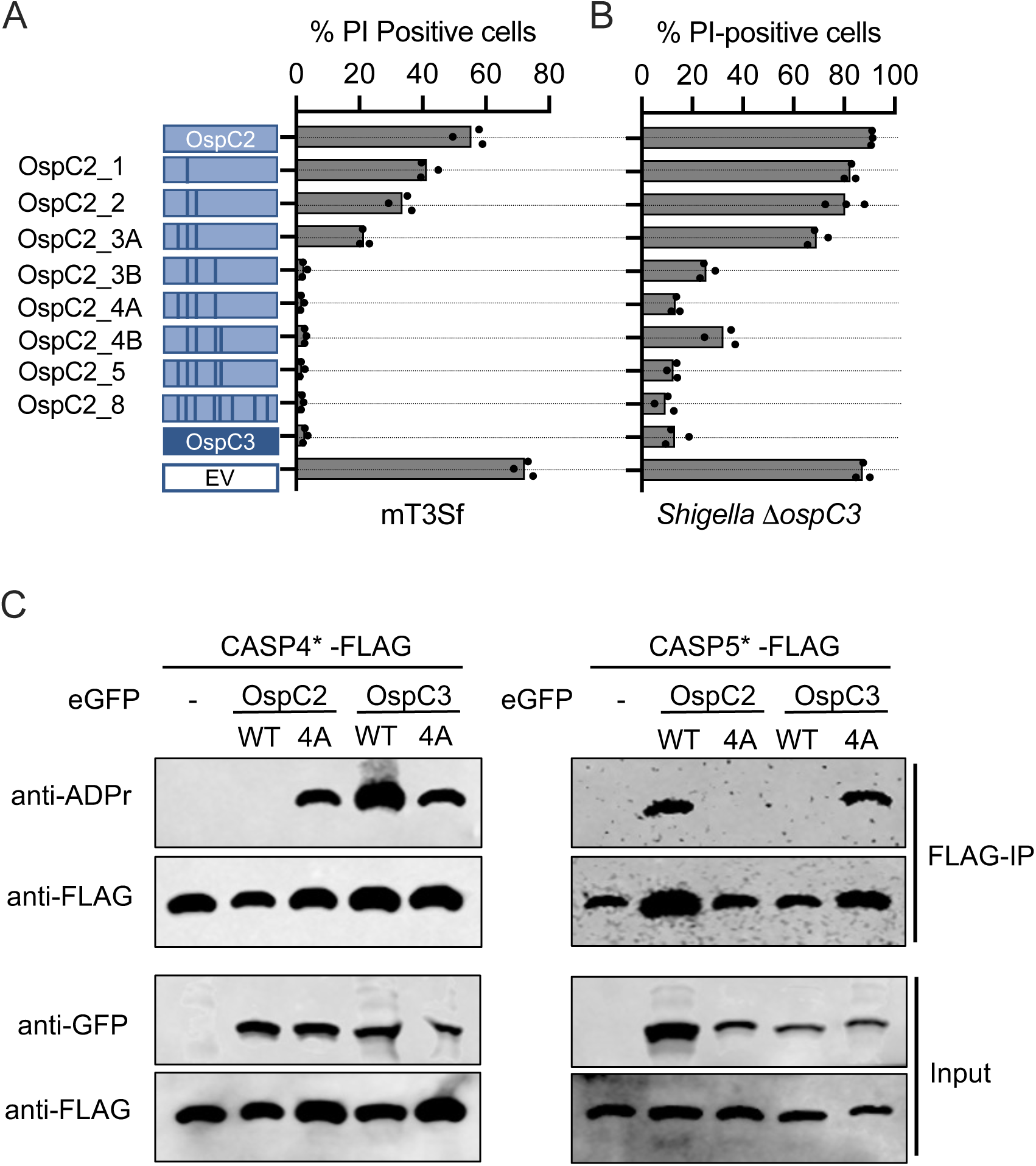
The PCS domains of OspC2 and OspC3 determine their binding specificity. (A, B) IFNγ-primed HeLa cells were infected with designated mT3Sf (MOI 5) (A) or Δ*ospC3* Shigella (MOI 3) strains (B). 1h post-infection with mT3Sf or 0.5h post-infection with Δ*ospC3 Shigella* strains, gentamicin, PI and Hoescht were added to the media, and the cells were imaged and the percentage of PI+ cells determined after an additional 2h. Each infection condition included at least three technical repeats and that shown is representative of 3 independent assays. (C) HEK293 cells were co-transfected with plasmids that constitutively express the designated GFP-OspC effector and catalytically dead full-length variant of CASP4 or CASP5. After 24h, the FLAG-tagged caspases were immunoprecipitated. Immunoblots of the input and immunoprecipitated fractions were probed with the designated antibodies are shown. The blots are representative of at least three experimental repeats.

Recognizing that neighboring residues could play a role in positioning or stabilizing the α-helix, we expanded the substitutions to include OspC3 residues at positions 423 and 425. Adding these additional residues led to the generation of variants OspC2_5, OspC2_4A, and OspC2_4B (Fig. 5A), each of which was as effective as OspC3 in suppressing mT3Sf-triggered cell death. Complementary changes in OspC3 resulted in variants whose addition to mT3Sf led to corresponding decreases in OspC3 meidated suppression of mT3Sf-triggered cell death (Fig. S7). All mutated OspC2 and OspC3 variants were secreted at levels comparable to that of WT OspC2 and OspC3 (Fig. S1B), confirming that differences in cell death suppression were not due to altered effector expression or secretion.

To further validate our model and confirm the specificity of OspC-mediated CASP4 inhibition, we evaluated the ability of the eight OspC2 variants to suppress Δ*ospC3 Shigella-*triggered cell death (Fig. 5B). In general, each OspC2 variant exhibited slightly reduced suppression of Δ*ospC3 Shigella* as compared to mT3Sf*-*triggered cell death. The smallest modification that fully suppressed Δ*ospC3 Shigella-*triggered cell death was OspC2_4A, a variant engineered with alterations at the three amino acids in the α-helix (407, 410, and 413), plus residue 423.

Based on these data and previous studies, we propose that two domains cooperate to promote the interactions of OspC2/OspC3 with CASP4/CASP5. The first domain is composed of the globular domains and intervening linkers of the ankyrin-rich repeats (ARRs), common to OspC1, OspC2, OspC3 and CopC, which, in the case of OspC3, were previously established to promote CASP4 binding. We propose that residues in this region promote general caspase binding. The second domain is the α-helix located upstream of the ARRs, which we designate as the <u>P</u>yroptotic <u>C</u>aspase <u>S</u>pecificity (PCS) domain. This domain is unique to OspC2 and OspC3 and determines their binding specificities for human pyroptotic caspases.

### OspC2 and OspC3 PCS domains determine which caspases they target

We suspect that we and others^10,17,19^ have not detected any evidence of Δ*ospC2 Shigella-*triggered cell death of epithelial cells, even of IFNγ-primed cells (Fig. S5B), due to the minimal levels of CASP5 expression in these cells^5^. To gain additional evidence that the PCS domains of OspC2 and OspC3 determine their caspase binding specificities, we compared the ability of the WT and PCS swapped variants of each (OspC2_4A and OspC3_4A) to modify CASP4 and CASP5 using the transient transfection assay described above. Notably, OspC3_4A, similar to OspC2, modified CASP5, whereas OspC2_4A, like OspC3, targeted CASP4 (Fig. 5C). OspC3_4A also displayed limited recognition of CASP4, presumably due to its overexpression in the transfected cells (Fig. 5C). These observations strongly support the PCS domains of OspC2 and OspC3 dictate their binding specificities for pyroptotic caspases.

### OspC2 likely binds to CASP5 amino acids under positive selection

Residues within mammalian proteins that confer protection against invading pathogens often undergo rapid evolution. This hallmark of positive selection is driven by the continuous arms race taking place between hosts and pathogens^42^. CASP4 and CASP5 originated from a gene duplication about 50 million years ago^37^, when they diverged from mouse CASP11. Intriguingly, CASP5, but not CASP4, possesses hallmarks of positive selection^37^. The majority of these rapidly changing residues in CASP5 map to its unique N-terminal extension; however, three are located in its L-subunit, in close proximity to the predicted OspC2 binding interface. Two of these positively selected residues in CASP5, positions 217 and 221 (Fig. 6A), are predicted by AlphaFold 3^43^ to be on an α-helix located in close proximity to the PCS domain of OspC2 (Fig. 6B). These two residues are among five in this 14 amino acid helix that differ between CASP4 and CASP5 (Fig. 6A).

**Fig. 6.**
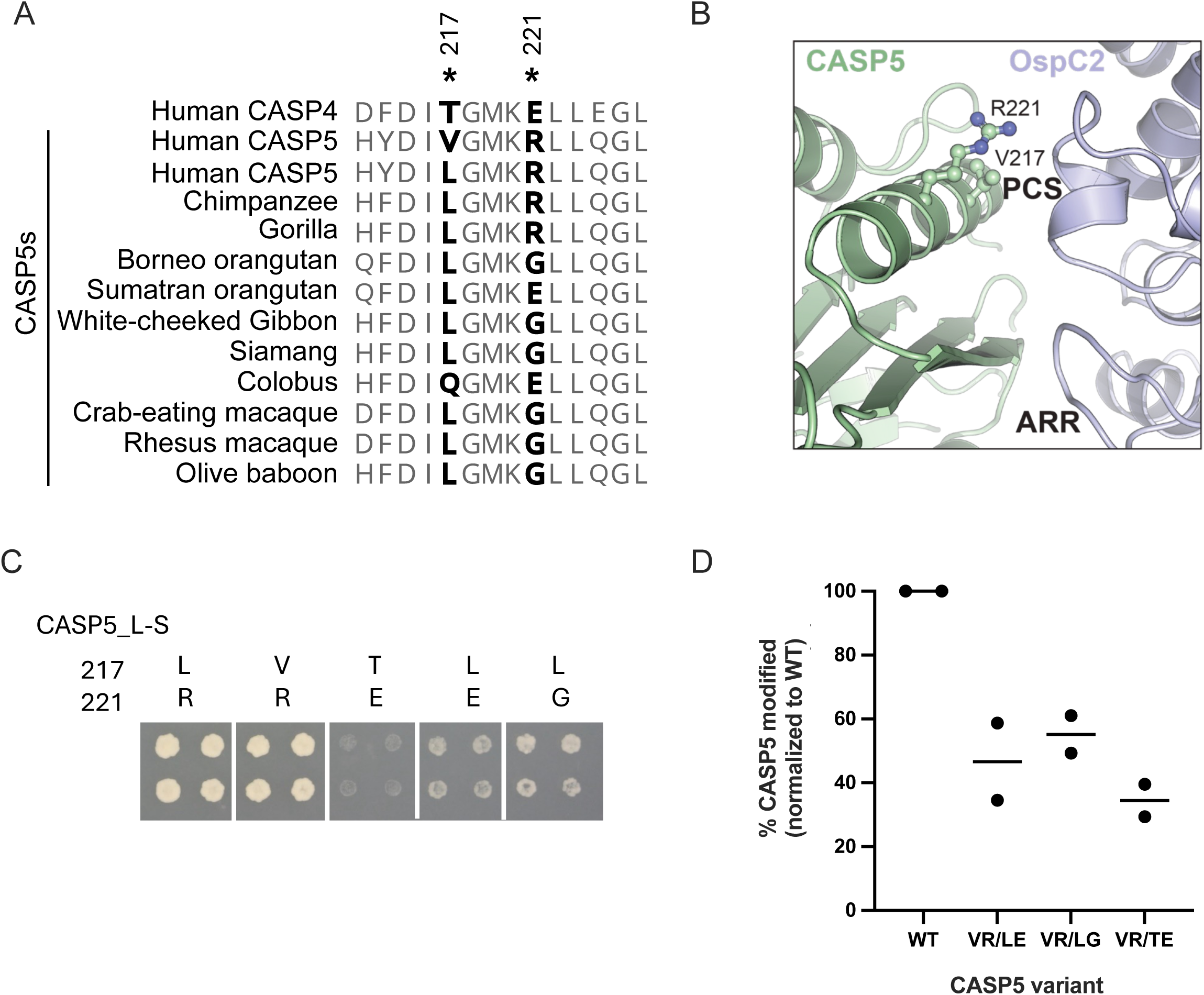
OspC2 likely binds to CASP5 amino acids that are under positive selection. (A) Sequence alignment of the residues of the outer α-helix on the L subunit of human CASP4/5 and CASP5 of Old-World primates. Positively selected CASP5 residues demarcated with asterisks. (B) Zoomed in region of AlphaFold 3 predicted complex of the L-S fragment of CASP5 in complex with the PCS and ARR domains of OspC2. (C) Y2H assay in which each quad spot represents yeast that co-express GAL4BD-OspC2 and the designated variant of GAL4AD-CASP5_L-S. (D) Quantification from two experiments of immunoprecipitated CASP5 modified by OspC2 in HEK293 cells transfected with plasmids encoding GFP-OspC2 and 3xFLAG-tagged, catalytically inactive CASP5 variants.

As a first test of whether these residues in CASP5 play a role in its binding to OspC2, we substituted them with the corresponding residues in CASP4. The resulting CASP5 variant no longer exhibited a Y2H interaction with OspC2 (Fig. 6C). Amino acid 217 in human CASP5 is usually a valine. However, in about 4% of people it is leucine or methionine, and leucine is found at this position of CASP5 in nearly all Old World primates (Fig. 6A)^37^. Substituting leucine 217 with valine had no effect in the Y2H assay, both variants show an equally strong interactions with OspC2 (Fig. 6C). Position 221 in human CASP5 is invariantly arginine, as it is in closely related non-human primates, i.e., chimpanzees and gorillas. In contrast, more distantly related orangutans have either a glycine (G) or a glutamic acid (E) at this position. Replacing R221 of human CASP5 with either a glycine or a glutamic acid results in a variant that barely interacts with OspC2 (Fig. 6C).

We next investigated how well variants of human CASP5 outfitted with the human CASP4 (TE) or orangutan residues (LE/LG) were modified when co-expressed with OspC2 in HEK cells. The immunoprecipitated CASP5-FLAG samples were immunoblotted for total CASP5-FLAG as well as modified CASP5. Band intensities on each blot were measured, and to account for experimental variability in transfection and immunoprecipitation efficiencies, the levels of the variant proteins on each immunoblot were normalized to those observed with the WT sample. The ratio of modified to total CASP5 was then calculated. Notably, OspC2 was less effective at modifying each CASP5 variant, with the greatest reduction observed with that containing the CASP4 substitutions at positions 217 and 221 (Fig. 6D, Fig. S8). Together these observations suggest a role for residues 217 and 221 of CASP5 in promoting its interaction with OspC2.

## Discussion

This study elucidates the distinct mechanisms by which *Shigella* OspC2 and OspC3 modulate human pyroptotic caspases, expanding our understanding of bacterial strategies for immune evasion in the intestinal epithelium. We found that despite sharing 96% identity, OspC2 and OspC3 uniquely modify human CASP5 and CASP4, respectively. Through a combination of Y2H, protein modeling, transient transfection, and infection studies with effector chimeras, we provide evidence that residues within their unique conserved PCS domains dictate the selective recognition and modification of CASP5 and CASP4 by OspC3 and OspC2, respectively. Reciprocal domain swaps between the two were sufficient to altered their caspase targeting profiles.

*Shigella* OspC2 and OspC3, like human CASP5 and CASP4, are closely related paralogs, each pair arising from a gene duplication event. Evolutionary pressures have driven the divergence of each pair. *Shigella* species are postulated to have appeared 270,000 years ago^44^, much later than the CASP4/CASP5 duplication event and the orangutan/gorilla split. The ancestral origin of the *Shigella* virulence plasmid, which encodes its T3SS and most of its effectors, including the OspC effectors, is unknown. We hypothesize that a bacterium carrying an effector with closely homology to OspC2 exerted selective pressures that led to the rapid evolution of CASP5 variants in primates. Furthermore, because the set of pathogens circulating in the wild among Old World primates is not fully characterized, there could even be *Shigella* species infecting these populations, perhaps with variants of OspC2 that have evolved to target the other CASP5 variants.

The physiologic significance of OspC2 targeting of CASP5 remains to be determined. Essentially nothing is currently known regarding the role of CASP5 in the defense of invading enteric pathogens, particularly within intestinal epithelial cells. This likely reflects, at least in part, its low expression in commonly studied intestinal cell lines^5^. Nevertheless, expression of CASP5, as well as CASP4, is up-regulated within colonic epithelial cells in states of inflammation in patients with inflammatory bowel diseases, suggesting that both are associated with colonic inflammation^45^. Thus, we hypothesize that via their complementary inhibition of pyroptosis, OspC2 and OspC3 promote the replication of *Shigella* within intestinal epithelial cells. Future studies will focus on investigating a role for *Shigella* OspC2-mediated suppression of CASP5-triggered pyroptosis using intestinal cell lines or colonic organoids under conditions identified to enhance CASP5 expression.

In sum, these findings highlight the evolutionary interplay between bacterial effectors and host proteins and suggest a role for CASP5 in the detection of *Shigella* and other Gram-negative bacteria. By elucidating the molecular basis of caspase targeting, this study advances our understanding of the diversification of host defense strategies. Furthermore, it identifies potential therapeutic targets to enhance epithelial cell resistance to bacterial infections and strengthen the host’s innate immune response, which could prove to be particularly useful in treating multidrug-resistant pathogens.

## Methods

Descriptions of the plasmids (and strains used in these studies are found in Tables S1-S3.

### Plasmid construction

Effector and caspase expression plasmids and were generated using a combination of Gateway recombination cloning, Gibson assembly, and site-directed mutagenesis, as summarized in Tables S1 and S2. For Gateway cloning, entry clones were generated by BP recombination into pDONR vectors and subsequently transferred into destination vectors by LR recombination (Invitrogen). Gibson assembly (NEBuilder HiFi DNA Assembly Kit, New England Biolabs) was used to insert PCR-amplified fragments or synthetic DNA into linearized backbones with 15–25 bp overlaps. When indicated, site-directed mutations were introduced by PCR-based mutagenesis (QuikChange II, Agilent) following the manufacturer’s instructions. All constructs were verified by Sanger sequencing.

### pmT3SS.1_LP construction

pmT3SS.1_LP was generated using a modified version of the strategy described by Reeves and Lesser^46^. Briefly, a DNA fragment that shares homology with *orfs131a* and *b* was PCR-amplified from the *Shigella* virulence plasmid and introduced in place of targeting sequence 2 (TS2) into pLLX13-*ipaJ-bla-spa40* via conventional cloning. The resulting new capture vector, pLLX13-*ipaJ-bla-orf131*, was digested with *MluI* and *PmeI* and introduced into VP_*E. coli* that carry pKD46 to generate pmT3SS.1 via homologous recombination. The resulting colonies were confirmed by colony PCR and next-generation sequencing. Once sequence verified, pmT3SS.1 and pNG162-VirB were introduced into DH10ß to generate mT3.1_*E. coli*.

### Strain construction

#### mT3Sf

mT3Sf was generated using the approach described in detail in Reeves et al^46^. First, using the αred recombination system, a synthetic 1.3 kb landing pad insertion site was introduced into the *atg/gid* locus of BS103 *Shigella* to generate BS103-LP^atp/gid^. The landing pad fragment plus flanking homology to the atg/gid locus via PCR amplification of pTKIP-tetA. Its flanking integration sites were confirmed by PCR. Next, pmT3SS3.1_LP, a plasmid that contains the Ipa, Mxi and Spa operons flanked by LP and *SceI* sequences was introduced into BS103-LPatp/gid + pKD46 via triparental mating: donor (DH10ß/pT3SS3.1_LP-Kan^R^), helper HB101 (pRK2073-spec^R^) and recipient (BS103-LPatp/gid/pKD46-Amp^R^). After curing the temperature sensitive pKD46, Spec^S^Tet^R^Kan^R^ BS103-LPatp/gid/pT3SS3.1_LP was transformed with pTKRED, which encodes the lambda recombinase and *SceI* restriction enzyme. The landing pad recombination system^47^ was then used to generate Kan^R^Tet^S^Spec^S^ mT3Sf candidates, which were screened for proper integration junctions by PCR. To generate mT3SfΔ*mxiE,* the Kan^R^ cassette present in mT3Sf adjacent to the Ipa operon was removed using pCP20. The αred recombination system was then used to remove most of *mxiE*, ensuring that the overlapping coding sequence of *mxiD* was maintained. The deletion was confirmed via PCR.

### *Shigella* deletion strain construction

Δ*ospC2 Shigella* were generated in *S. flexneri* 2457T via the αred recombination system^48^.

### Y2H assay

The Y2H expression plasmids were introduced into MaV103 and MaV203, and the Y2H assays were performed in a 96-well format as previously described^49^. In this case, the two-hybrid selection was conducted on medium lacking leucine, tryptophan and histidine, supplemented with 40 mM 3-amino-1,2,4-triazole. Growth was scored after 7d incubation at 30°C.

### Liquid secretion assay

Liquid Secretion assays were performed as previously described^21^ with some modifications. Overnight cultures of mT3Sf or *Shigella* were grown at 30°C in TCS. Antibiotics were added for mT3Sf for plasmid maintenance. In AM, cultures were diluted 1:100 into 2.5 mLs of TCS and grown with shaking at 37°C. For mT3Sf, 1 mM IPTG was added 1h into the incubation. Once cultures reached OD600 ∼1.0, equivalent numbers of bacteria were pelleted, resuspended in 2.5 mL PBS containing 10 µM Congo red and incubated with shaking for 0.5h at 37°C. Following the incubation, two consecutive centrifugations were used to separate pellets and supernatant fractions. Proteins in the supernatant fractions were TCA precipitated before being suspended into loading dye. Cell equivalent amounts of supernatant and whole cell lysate samples were run on an SDS-PAGE 12% Tris-glycine gel, transferred to a nitrocellulose membrane, and probed with the indicated antibodies: anti-FLAG (1:5000), anti-GroEL (1:100,000, Sigma, G6532), anti-IpaB (1:20,000), anti-IpaC (1:40,000), anti-IpaD (1:40,000), anti-RNA Pol (1:5,000). Western blots were visualized using an Azure 600 imager.

### Solid plate secretion assay

Solid plate secretion assays were conducted as previously described^50^. Single wells of a 96-well plate were inoculated with mT3Sf that carry a plasmid expressing designated effector. The plate was incubated with agitation overnight at 37°C. The following day, a BM3-BC pinning robot (S&P Robotics Inc., Toronto, Canada) was used to quad-spot equivalent volumes of each culture onto solid TCS agar trays containing 10 µM Congo red and 1 mM IPTG agar. Immediately after transfer, a nitrocellulose membrane was overlaid. After a 4.5h incubation at 37°C, the membrane was removed, washed three times with TBS-T (Tris-buffered saline with 0.1% Tween-20) and then immunoblotted with the M2 α-FLAG (1:5000) antibody (Sigma) before been visualized using an Azure 600 imager.

### Mammalian cell lines

HeLa, HCT8, and HEK293 cells obtained from the American Type Culture Collection were cultured in a 5% CO2 incubator at 37°C. HeLa and HEK293 cells were maintained in high glucose DMEM (Thermo Fisher Scientific) and HCT8 cells in GlutaMAX-supplemented RPMI 1600 (Thermo Fisher Scientific). In each case, media were supplemented with 10% heat-inactivated fetal bovine serum (FBS, R&D systems), 100 IU/mL penicillin, and 100 µg/mL streptomycin) (Life Technologies).

### Knockout cell lines

CASP4 knockout cells were generated via CRISPR/Cas9 editing. The sgCASP4 (CTTCATGAGGACAAAGCTTG) was mixed with SpCas9 2NLS nuclease (Synthego) to generate ribonucleoprotein (RNP) complex. The RNP complexes were electroporated into 1x10^6^ cells (HeLa) using a Neon system (Thermo Fisher Scientific). Pools were monitored for knockout efficiency after a 2–3 day recovery. Cells were diluted to isolate single cells, which were expanded to generate clonal cell lines. Knockouts were verified by immunoblotting lysates with anti-CASP4 antibody (Santa Cruz, clone 4B9, sc-56056).

### Gentamicin/Chloroquine protection assay

HeLa cells seeded in 96-well plates (2 × 10^4^ cells per well) were infected with *Shigella* (MOI 100), mT3Sf (MOI 10) or *E. coli* expressing *Yersinia* Invasin (MOI 10). After 1h, gent (100 μg/mL) ± CHQ (200 µg/mL, 400 µM) was added to each well for 1h. The infected cells were then washed three times with PBS and then lysed with PBS containing 1% (vol/vol) Triton X-100. Serial dilutions of the lysates were plated, and the number of bacteria were enumerated. Percent cytosolic bacteria were determined by calculating the ratio of (CHQ^R^ + Gent^R^)/Gent^R^ bacteria. Percentages > 100% were capped at 100%.

### Infection and cell death assay

One day prior to infection, mammalian cells were seeded in antibiotic-free media at 2x10^4^ into 96-well tissue culture treated plate (Corning). For IFNγ-priming, the cells were treated with 10 ng/ml IFNγ overnight (0.1% BSA, Peprotech) when seeded. Just prior to infection, the media was replaced with low-glucose DMEM. For mT3Sf infections, single colonies were picked and grown overnight in 2 mL TCS broth at 37°C with aeration. For *Shigella* infections, single Congo red colonies were picked and grown overnight in 2 mL TCS (trypticase soy) broth at 30°C with aeration. The next day, each culture was diluted 1:50 into TCS broth and grown at 37°C with aeration until an OD600 of 0.6 to 1.0 (2 h approximately). For mT3Sf, 1mM IPTG was added 1h post back-dilution. Bacteria resuspended in prewarmed low-glucose DMEM (no phenol red) supplemented with 1% FBS were added to HeLa cells in 96-well plates at designated MOI after which plates were centrifuged at 2,000 rpm for 10 min to synchronize the infection. For mT3Sf infection, IPTG (1mM) was maintain throughout the infection. All strains used for these studies expressed AfaI^51^. The plates were incubated at 37°C in a humidified 5% CO2 incubator for 1h of mT3Sf and 0.5h for *Shigella*, after which each well was washed two times and then incubated in Plate Reader Media containing HBSS (Hanks’ Buffered Salt Solution), 10% FBS, 50 mM HEPES plus gentamicin (50 µg/mL), 3 µM PI and 16.2 µM Hoechst 33342. The plate was imaged using an ImageXpressPICO automated microscope (Molecular Devices) using a 20x/0.4 lens with DAPI and Texas Red filters. Images were obtained every 10-15 min. Images of each well were analyzed using the Cell Reporter Xpress software (Molecular Devices). The percentage of dead cells was calculated by dividing the total number of PI-positive cells by the total number of Hoechst-stained cells imaged in each well. In each case, the results from three replicate wells were averaged.

### Transfection and immunoprecipitation assay

To assess ADP-riboxanation of total cell proteins by OspC variants, HEK293T cells seeded in 6 well tissue culture plates (CellTreat) were transfected with 300 ng of DNA using Fugene6 (Promega). After 24h, the cells were lysed in 250 µL ice-cold radioimmunoprecipitation assay (RIPA) buffer (25 mM Tris, pH 8, 150 mM NaCl, 0.1% SDS, and 1% NP-40 plus mini-EDTA free protease inhibitor cocktail; Millipore Sigma). After a 15 minutes incubation in RIPA buffer, the soluble fractions were obtained by centrifugation at 13,200 g for 10 min at 4°C. The pellet was discarded and supernatant denatured in SDS loading buffer and loaded onto a 12% SDS-PAGE gel, followed by immunoblotting with anti-ADP ribose (Clone D9P7Z, #89190, Cell Signaling) and anti-HSP90-HRP conjugate (Clone C45G5, #79641, Cell Signaling) antibodies.

To assess ADP-riboxanation of FLAG-tagged caspases by OspC variants, 2.4x10^6^ HEK293T cells were seeded in 100mm x 20mm tissue culture treated dishes (CellTreat). The cells were co-transfected with 850 ng of the appropriated GFP-OspC and FLAG-tagged caspase (5.6 µg of caspase-4 or 15 µg of caspase-5) expression plasmids using Fugene6. After 24h, the cells were lysed in 300 µL of ice-cold lysis buffer (50 mM Tris-HCl (pH 7.6), 150 mM NaCl, 5mM MgCl2, 2 mM EDTA, 100 µg/mL Digitonin plus mini-EDTA free protease inhibitor cocktail). The concentration of total proteins in each lysate was determined using Bradford assay. Anti-Flag M2 magnetic beads (M8823 Millipore Sigma), pre-washed with IP buffer (50 mM Tris-HCl (pH 7.6), 150 mM NaCl, 5mM MgCl2, 2 mM EDTA, 20 µg/mL Digitonin) were added to 1-2 mg/ml soluble fraction and incubated for 2h at 4°C with constant rotation. The beads were then washed five times with the IP buffer and then eluted with SDS loading buffer. Samples were loaded onto a 12% SDS-PAGE gel, followed by immunoblotting with anti-Flag, anti-GFP (66002-1, Proteintech) and anti-ADP-ribose antibodies, using conditions described above.

### Sequence and structural analyses

The AlphaFold3 web interface was used for all structural modeling^43^. Model confidence was assessed using pTM and ipTM scores, as well as PAE plots as described. Structural analysis was performed using PyMol (The PyMOL Molecular Graphics System, Version 3.0 Schrödinger, LLC). Sequence alignments of OspC homologues were generated with T-Coffee and visualized using JalView^52^.

### Statistical analyses

Prism 10 (GraphPad Software) was used to graph data and conduct all statistical analyses. Differences were considered statistically significant when P value was <0.05.

## Acknowledgements

We thank Drs. Patrick Mitchell, Isabella Rauch and Charlotte Odendall for critically reading the manuscript and members of the Vance, Odendall, Mitichell, Goldberg and Barczak lab members for helpful discussions. We thank Analise Reeves, Sarah Bier and Mikaela Svensson for helping generate plasmids and strains and Dr. Sunny Shin (University of Pennsylvania) for sharing reagents. This work was supported by NIH grants AI169795 to C.F.L, AI007061 to K.K., R35GM150681 to TCL and R35GM142486 to JNP.

## References cited in Supplemental Tables (not cited in main text)

^53 54 55 56 57 58 59 60 61^

