## Supplemental Figures for "Differential targeting of human pyroptotic Caspase-5 and Caspase-4 by *Shigella* OspC2 and OspC3"

Figure S1

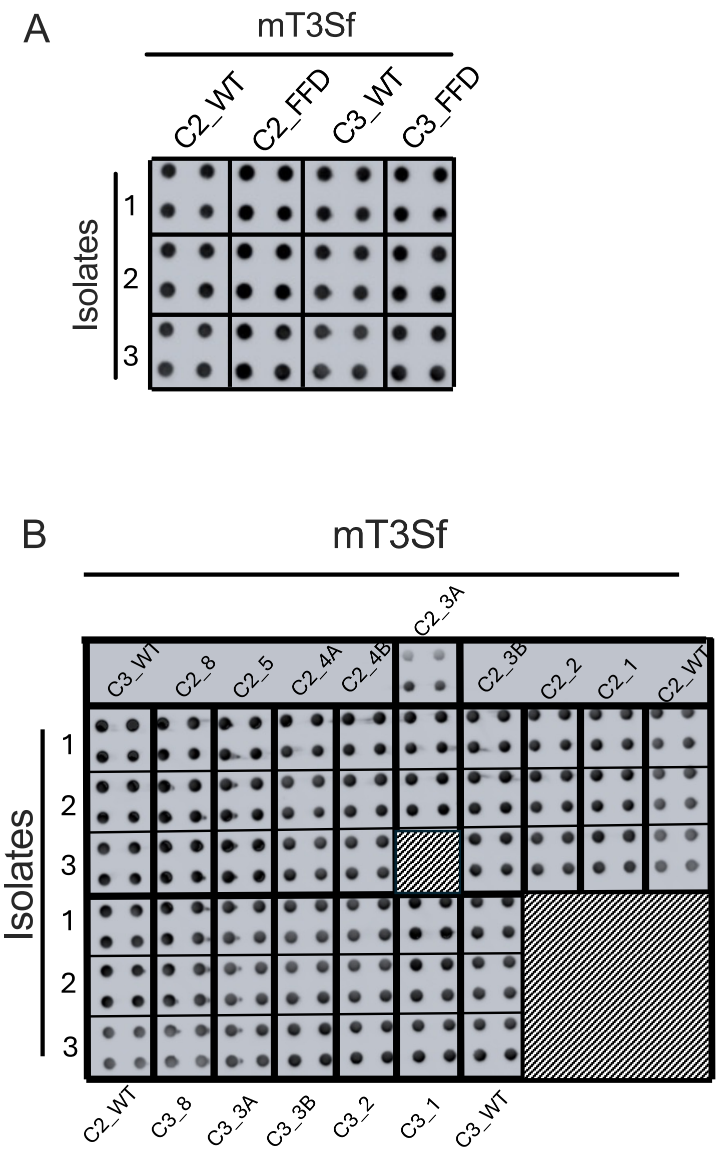


**Fig. S1 OspC variants are secreted at levels equivalent to WT effectors.** (A-B) Solid plate secretion assay of mT3Sf expressing designated OspC variants. Blots were probed with anti-FLAG antibody. Each set of four spots originated from a single isolate.

Figure S2

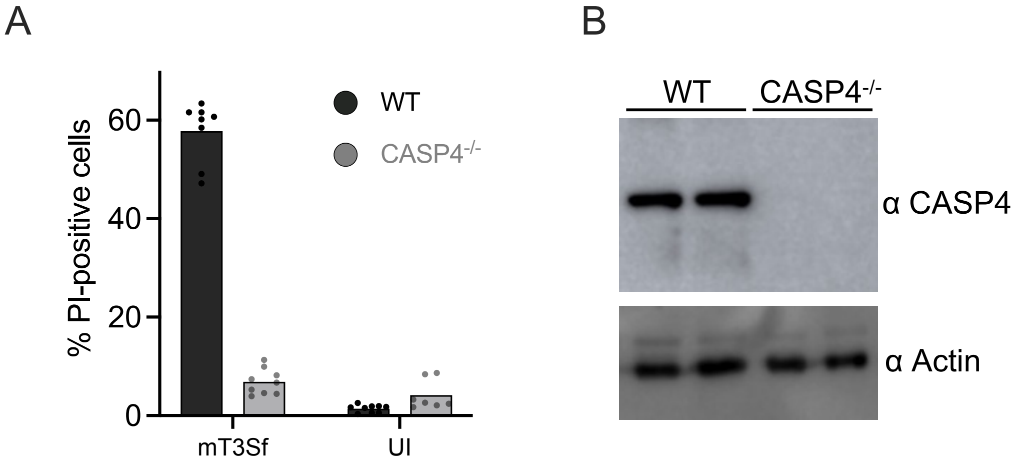


**Fig. S2 Infection with mT3Sf triggers CASP4-dependent pyroptosis of epithelial cells.** (A) IFNγ-primed WT or CASP4$^{-/-}$ HeLa cells were infected with mT3Sf at an MOI of 5. One-hour later, gentamicin, PI and Hoechst were added to the media. After Two hours, the cells were imaged and the percentage of PI+ cells quantified. Each infection condition included at least three technical repeats and that shown is representative of at least 3 independent assays. (B) Immunoblots of whole cell lysates two independent CASP4$^{-/-}$ HeLa cells probed with designated antibodies.

Figure S3

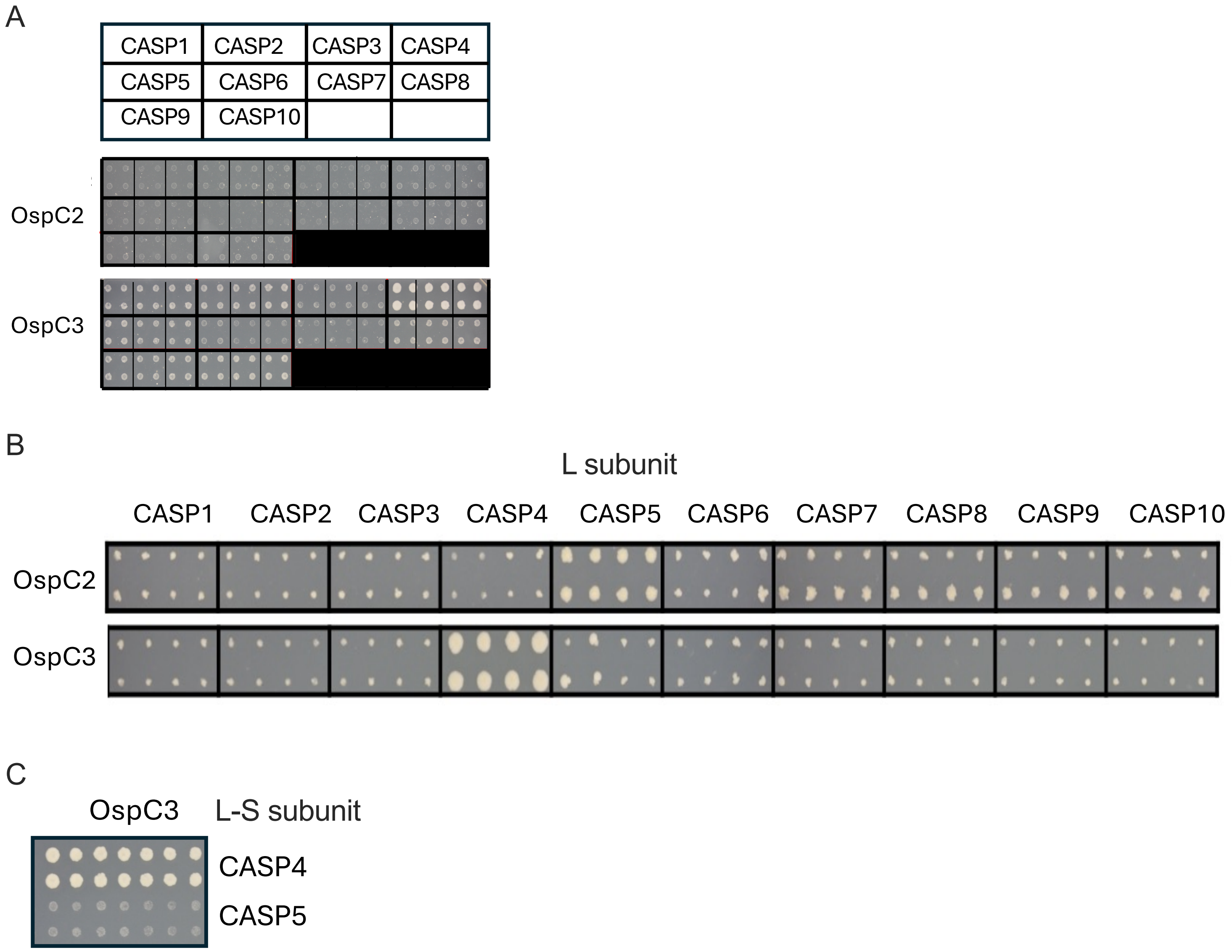


**Fig. S3 OspC2 and OspC3 specifically interact with and target CASP5 and CASP4, respectively.** (A-C) Y2H assays were performed to assess the interactions of OspC2 and OspC3 with full-length human caspases 1-10 (A), their respective L domains (B), and OspC3 with the L-S domain of CASP5 (C). Images shown were obtained after seven days of incubation on selective media.

Figure S4

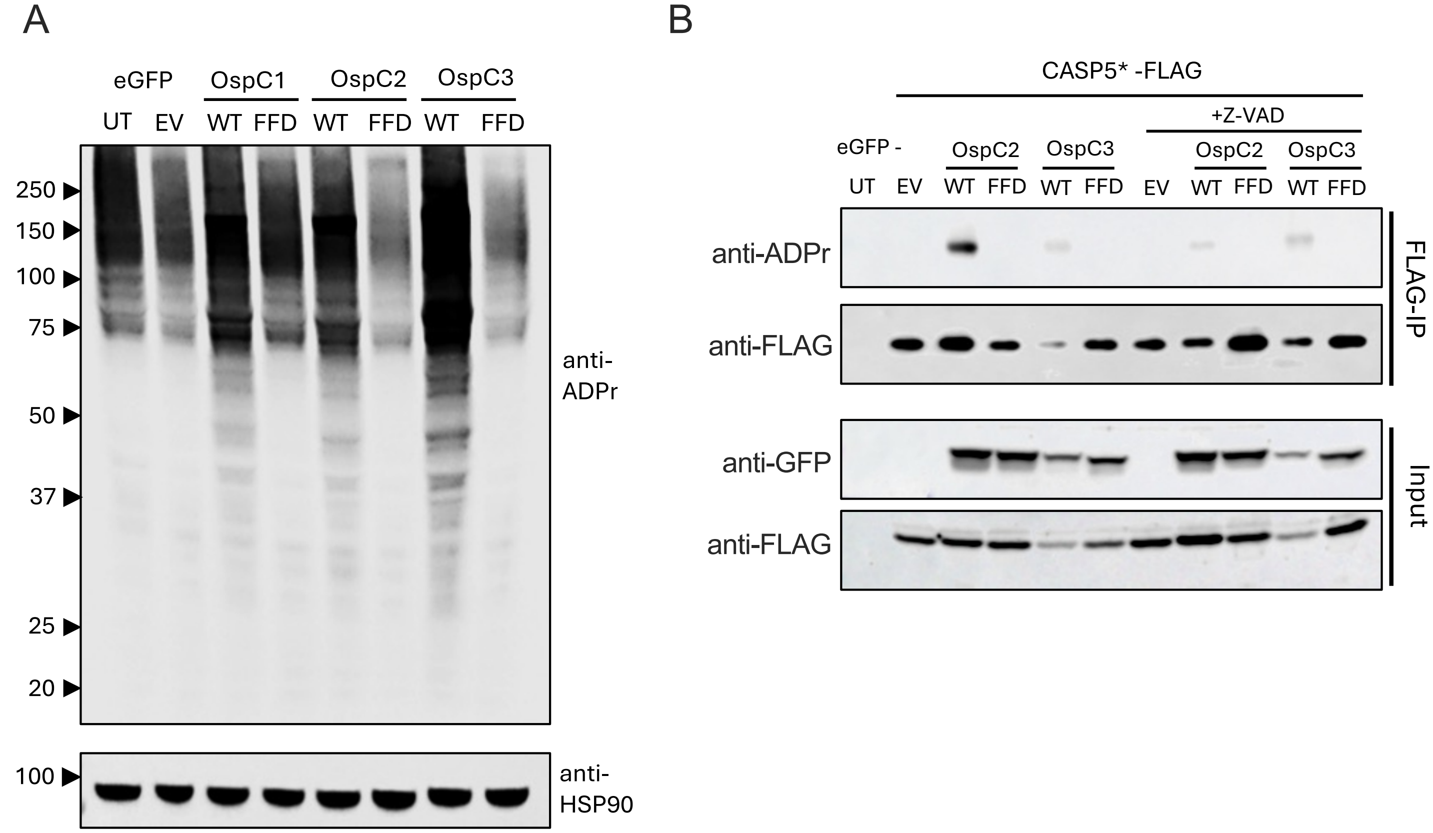


**Fig. S4 ADP-riboxanation profiling of OspC variants.** (A) HEK293 cells were transfected with a plasmid that expresses a WT or catalytically dead GFP-OspC variant. After 48h, cells were lysed and the whole cell lysates were probed with the designated antibodies. (B) HEK293 cells were co-transfected with plasmids that express the designated GFP-OspC variant and one the expresses 3xFLAG tagged catalytically dead CASP5 (B). After 24h, ZVAD was added to the media. After an additional 48h the cells were lysed, and the FLAG-tagged caspases were immunoprecipitated. Immunoblots of the input and immunoprecipitated fractions were probed with the designated antibodies. Panel (A) shows results from three experiments; panel (B) reports ZVAD effect from one experiment.

Figure S5

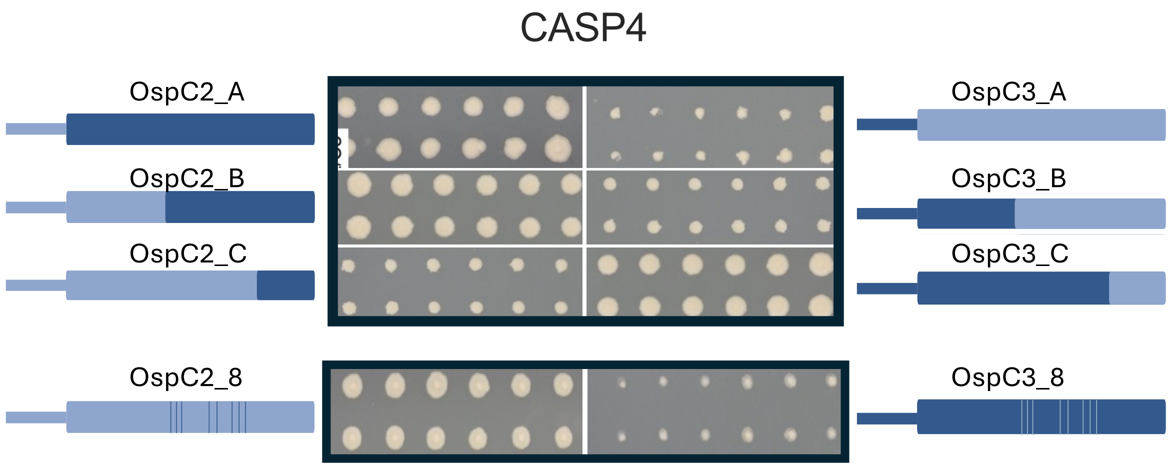


**Fig. S5 Identification of OspC chimeras that bind to full-length CASP4.** (A) Y2H assays were performed to assess the interactions between OspC chimeras and full-length CASP4. Images obtained after 7 days of growth on selective media.

Figure S6

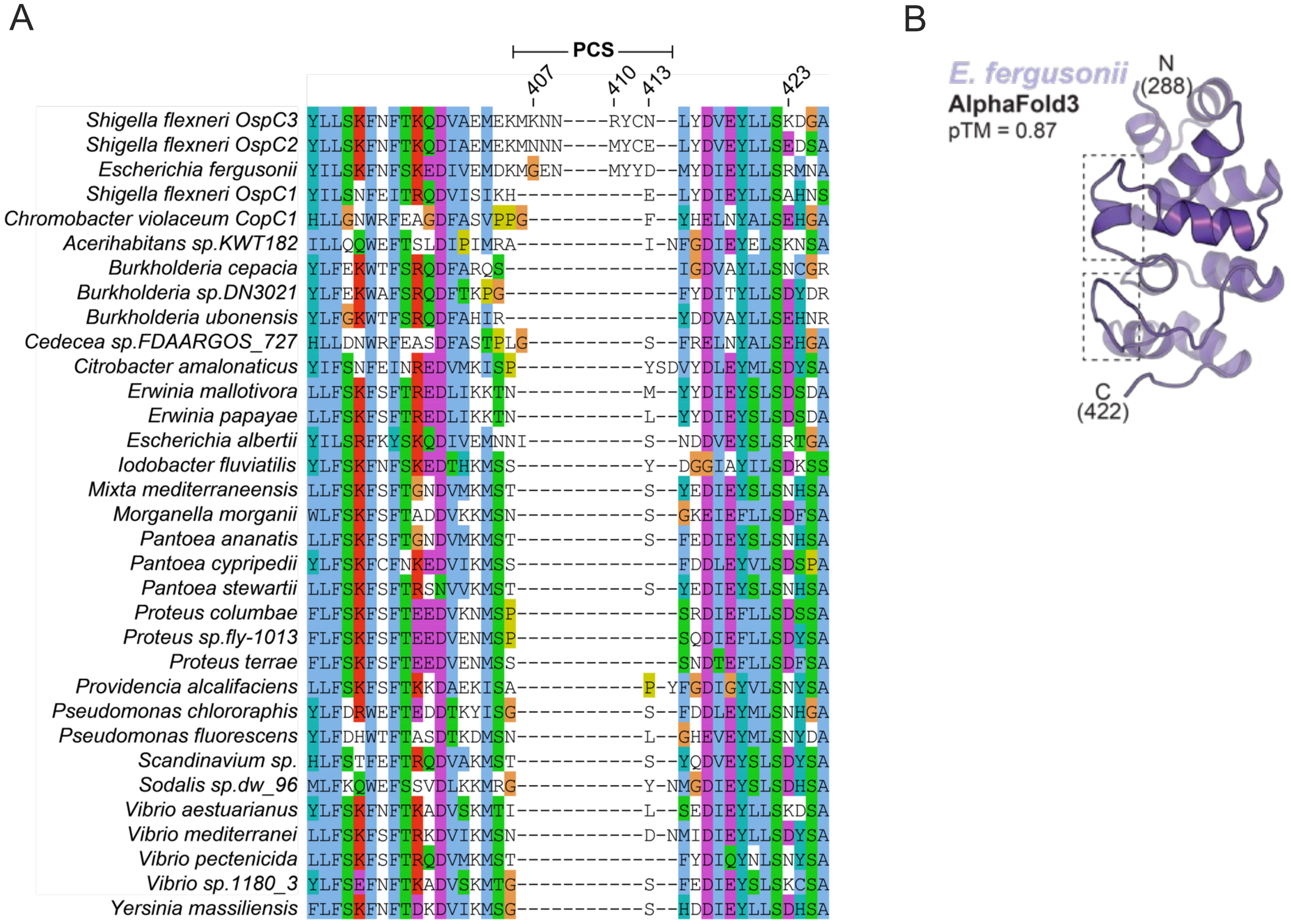


**Fig. S6 Alignment of ARDs of OspC homologs.** (A) Alignment of upstream region of ARD domains of OspC homologs identified via a BLAST search with full length OspC3. (B) AlphaFold3 prediction of the structure of the ARD of *E. fergusonii.*

Figure S7

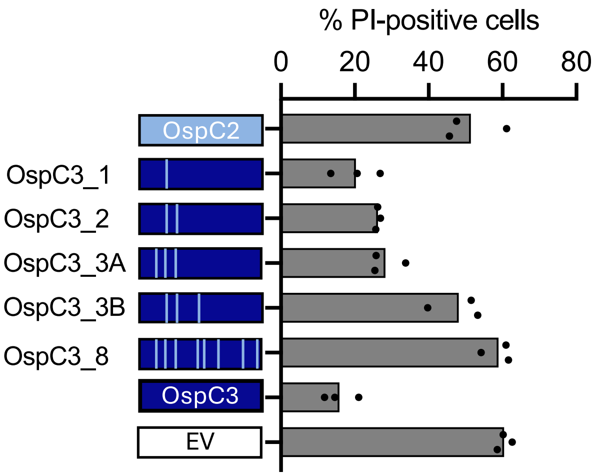


**Fig. S7 Characterization of the ability of OspC3 variants to suppress mT3Sf-triggered cell death** IFNγ-primed HeLa cells were infected with designated mT3Sf strain at MOI of 5. One-hour post-infection, gentamicin, PI and Hoescht were added to the media. After an additional 2h, the cells were imaged and the percentage of PI+ cells determined. Each infection condition included at least three technical repeats and that shown is representative of at least 3 independent assays.

Figure S8

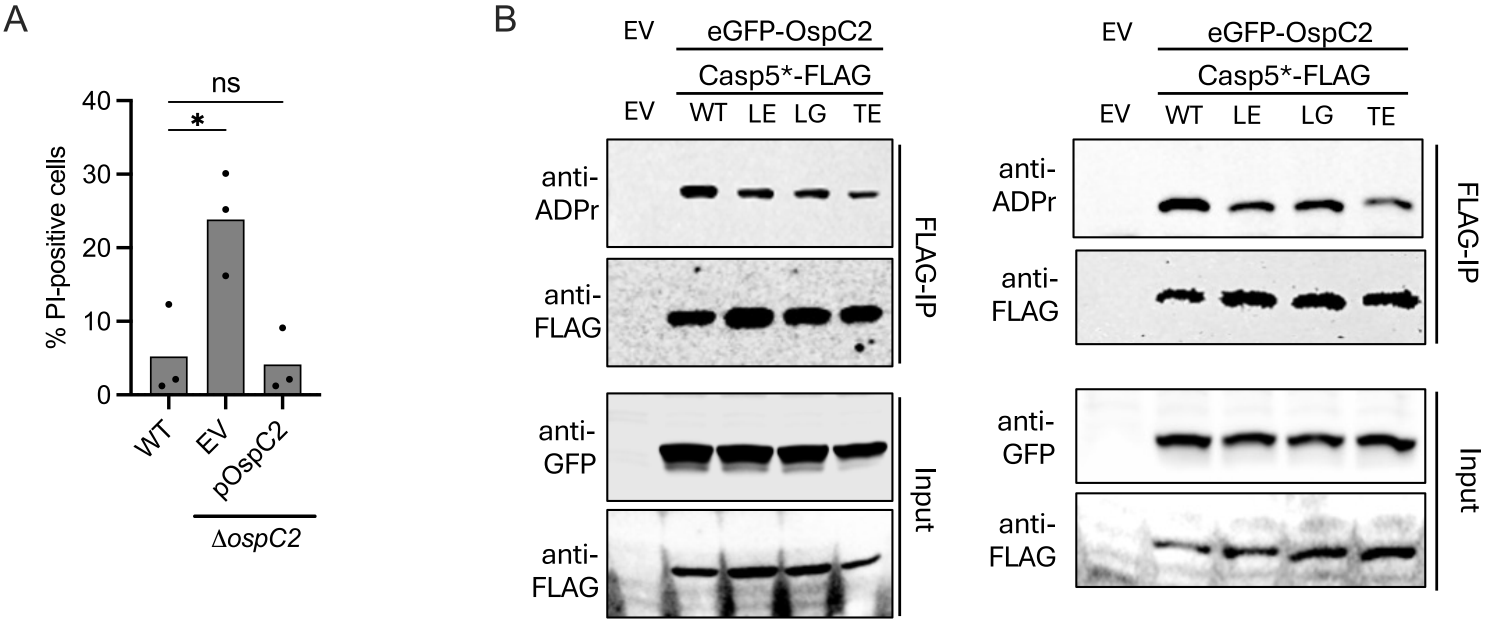


**Fig. S8 OspC2 modifies CASP5 through recognition of positively selected residues in an exposed CASP5 α-helix.** (A) IFNγ-primed HeLa cells were infected with designated WT, Δ*ospC3* or Δ*ospC3 Shigella* at an MOI 10. Thirty minutes post-infection gentamicin, PI and Hoescht were added to the media. After an additional 2h, the cells were imaged and the percentage of PI+ cells determined. Statistical significance when indicated was assessed by one-way ANOVA with Tukey's post hoc test. *P < 0.05, **P < 0.01, ****P < 0.0001, ns = nonsignificant. (B) HEK293 cells were co-transfected with plasmids that express GFP-OspC2 and the indicated 3xFLAG tagged catalytically dead full-length variant of CASP5. After 24h, the cells were lysed and the FLAG-tagged CASP5s were immunoprecipitated. Immunoblots of the input and immunoprecipitated fractions were probed with the designated antibodies. The blots shown are representative of two experimental repeats.
